# Improved Docking of Protein Models by a Combination of Alphafold2 and ClusPro

**DOI:** 10.1101/2021.09.07.459290

**Authors:** Usman Ghani, Israel Desta, Akhil Jindal, Omeir Khan, George Jones, Nasser Hashemi, Sergey Kotelnikov, Dzmitry Padhorny, Sandor Vajda, Dima Kozakov

## Abstract

It has been demonstrated earlier that the neural network based program AlphaFold2 can be used to dock proteins given the two sequences separated by a gap as the input. The protocol presented here combines AlphaFold2 with the physics based docking program ClusPro. The monomers of the model generated by AlphaFold2 are separated, re-docked using ClusPro, and the resulting 10 models are refined by AlphaFold2. Finally, the five original AlphaFold2 models are added to the 10 AlphaFold2 refined ClusPro models, and the 15 models are ranked by their predicted aligned error (PAE) values obtained by AlphaFold2. The protocol is applied to two benchmark sets of complexes, the first based on the established protein-protein docking benchmark, and the second consisting of only structures released after May 2018, the cut-off date for training AlphaFold2. It is shown that the quality of the initial AlphaFold2 models improves with each additional step of the protocol. In particular, adding the AlphaFold2 refined ClusPro models to the AlphaFold2 models increases the success rate by 23% in the top 5 predictions, whereas considering the 10 models obtained by the combined protocol increases the success rate to close to 40%. The improvement is similar for the second benchmark that includes only complexes distinct from the proteins used for training the neural network.

## 1. Introduction

In Round 14 of Critical Assessment of Structure Prediction (CASP14), the doubly blind, biennial experiment focusing on protein structure prediction, the Google affiliated company DeepMind surprised the organizers by demonstrating major improvement in prediction accuracy [1]. The models had been obtained by AlphaFold2, a neural network based program. The AlphaFold2 algorithm and program has been recently released [2], and it is now generally recognized in the scientific community that DeepMind essentially solved the protein folding problem [3]. Given only the sequence, for a large fraction of monomeric proteins, the method is able to produce models that are indistinguishable from experimental structures. Although the accuracy may depend on the depth of the multiple sequence alignment (MSA) available for a given target, results are generally very good even without any meaningful templates and only with a shallow MSA.

Given the success of AlphaFold2 for protein structure prediction, the obvious question is whether the method can also predict protein-protein complexes. DeepMind did not participate in the assembly prediction experiment of CASP14 [4], but results show that most residues located at domain-domain interfaces of multi-domain proteins were predicted remarkably well. In fact, some of the predicted residue conformations seemed well placed for the interaction, in spite of not modeling the interacting protein partner. It was shown that docking models of interacting proteins generated by AlphaFold2 provides results that are comparable to docking X-ray structures [5]. Therefore, it was expected that deep neural network based methods can also be directly used for predicting the structures of protein-protein complexes. This assumption was confirmed by the Baker group by using RoseTTaFold, a different but similar deep learning based structure prediction method [6]. Rather than providing the sequence of a single protein as the input, they used two or more sequences with a gap between them, and demonstrated that the program can generate coordinates of two or more interacting protein chains. This was done by providing a paired alignment and modifying the residue index to include the gap. Thus, the network enabled the direct building of structure models for protein-protein complexes from sequence information, short circuiting the standard procedure of building models for individual subunits and then carrying out rigid-body docking. Mirdita et al. [7] has shown that the same technique works with AlphaFold2, often even without the paired alignment. AlphaFold2 uses relative positional encoding with a cap at |i – j| ≥ 32, which means that by offsetting the residue index between two proteins to be >32, AlphaFold2 treats them as separate polypeptide chains.

A similar protocol was applied to protein-peptide docking by Ko and Lee [8] and by Tsaban et al. [9]. Both groups linked the peptide sequence to the protein sequence via a polyglycine linker, and used AlphaFold2 without any modifications. The method was tested on large benchmark sets of protein-peptide complexes. There were some differences in the overall quality reported, but both studies concluded that fairly accurate docked structures could be obtained for about half of the targets. In some cases, the method obviously failed with the poly-glycine linker throwing the peptide segment into space. In other cases the models correctly identified the binding pocket, but showed errors of peptide rotation or translation [9]. Tsaban et al. also compared the performance of AlphaFold2 to that of the physics-based peptide docking protocol PIPER-FlexPepDock [10] and observed an almost orthogonal behavior in terms of successes and failures.

In the present paper we describe a protocol for protein-protein docking that substantially improves the AlphaFold2 generated models by combining AlphaFold2 with our established physics-based docking method ClusPro [11]. In the first step we use AlphaFold2 as described previously, with the two protein sequences separated by a gap, and generate five primary AlphaFold2 models. However, we go beyond what has been done in earlier studies and combine AlphaFold2 with ClusPro at three different points of the algorithm. First, the best AlphaFold2 model is selected, the two proteins in the complex are separated, and docked to each other using ClusPro. As standard with the server, ClusPro generates 10 models [11]. Each of the 10 ClusPro models are then used as templates in AlphaFold2 runs, resulting in 10 secondary AlphaFold2 models. As the third combination of the two methods, the five primary AlphaFold2 models from the first step are added to the 10 secondary models, and all 15 models are ranked using the PAE (Predicted Aligned Error) values generated by Alphafold2, selecting the model with the lowest PAE as the top prediction. As will be described, we tested the method on two benchmark sets of interacting proteins, and observed substantial increase in the number of complexes that were modeled with acceptable or better accuracy.

## 2. Methods

### 2.1. Benchmark sets

The combined method has been tested on two benchmark sets. Benchmark 1 was based on Version 5 of the very established protein-protein docking benchmark (BM5) developed by the Weng group [12]. BM5 is a nonredundant and fairly diverse set of protein complexes for testing protein-protein docking algorithms. Each entry in BM5 includes the 3D structures of the complex and one or both unbound component proteins. The set includes 40 antibody-antigen, 88 enzyme-containing and 102 “other type” complexes [12]. While Alphafold was applied to predict all 230 targets in BM5, due to the limitations of our GPU resources we were able to get results only for 204 complexes, among them were 34 antibody-antigen pairs and 170 nonantibody containing complexes that included both enzyme-containing and “other type” structures [12]. A drawback of using Benchmark 1 is that the proteins forming the complexes in BM5 were present in the training set used by AlphaFold2, which was trained on Protein Data Bank (PDB) structures released before May 2018 [2]. Thus, while the training set included only monomeric proteins, testing the method on BM5 may introduce some bias, leading to overestimating the performance of AlphaFold2.

Benchmark 2 was designed to exclude the possibility of implicit bias. It includes only heterodimeric targets from the PDB released after May 2018, since such protein structures were certainly not used for training AlphaFold2. Furthermore, we used ClusPro TBM to look for homologous templates for potential targets and limited our selection to those with no appropriate template in the PDB [13]. This was done to ensure that our data set consisted of truly novel protein-protein interactions. The final Benchmark 2 consists of 17 complexes with the PDB IDs 5ZNG, 6A6I, 6GS2, 6H4B, 6IF2, 6II6, 6ONO, 6PNQ, 6Q76, 6U08, 6ZBK, 7AYE, 7D2T, 7M5F, 7N10, 7NLJ, 7P8K.

### 2.2. Docking protein pairs using AlphaFold2

For the complexes in Benchmark 1, the sequences for the unbound proteins were extracted from BM5 in fasta file format [12]. For the targets in Benchmark 2, the sequence and chain information was obtained from the PDB using the PISA/author assigned biological assembly. We then generated a multiple sequence alignment (MSA) for each sequence using the MMseqs2 API [7, 14]. We joined the MSAs to make pseudo-monomers in which each chain of the oligomer was separated by gaps that were set to 200 residues in our protocol [7]. The final MSA was then used as the input without any template to build five AlphaFold2 models with the pTM model parameter set as this allowed us to also obtain the predicted aligned error (PAE) values for each model [2].

### 2.3. Calculating the average interface PAE

The PAE values calculated by AlphaFold2 provided an error estimate for each residue with respect to every other residue in the model. We used this to calculate an average interface PAE score, where interface was defined as the residues within 10.0 Å of the other subunit. For each residue on the interface, we retained the PAE values for the interface residues on the other subunit. These were then used to calculate the average interface PAE scores that provided the ranking of the models, with lower scores being ranked higher.

### 2.4. Docking predicted subunits using ClusPro

The residues in the top ranked model predicted by AlphaFold2 were split into two groups. One served as the receptor, and the other, the ligand. For the complexes in Benchmark 1, this split was based on the sequences of the two proteins as defined in BM5. The residues belonging to the first protein, usually the larger one, were defined as the receptor, and the residues belonging to second protein formed the ligand. For the heterodimeric targets in Benchmark 2, the sequences in the PDB files were used to define the two proteins. The separated receptor and ligand models were then submitted to the ClusPro server for docking [11]. Some chains had long unstructured tails which were manually cut before submission. The antibody-antigen pairs were docked using ClusPro’s antibody mode, whereas for all other targets we used the electrostatic-favored coefficient set as they have been shown to produce the best results [15]. In both cases the 10 best models generated by ClusPro were retained.

The ClusPro free docking protocol consists of two main steps [11]. The first step is running PIPER, a docking program that performs systematic search of complex conformations on a grid using the fast Fourier transform (FFT) correlation approach [16]. The scoring function includes the van der Waals interaction energy, an electrostatic energy term, and desolvation contributions calculated by a structure based pairwise potential [11]. The second step of ClusPro is clustering the top 1000 structures generated by PIPER using the pairwise RMSD as the distance measure. The radius used in clustering is defined in terms of C_α_ interface RMSD. For each docked conformation, we select the residues of the ligand that have any atom within 10 Å of any receptor atom, and calculate the C_α_ RMSD for these residues from the same residues in all other 999 ligands. Thus, clustering 1000 docked conformations involves computing a 1000 × 1000 matrix of pairwise C_α_ RMSD values. Based on the number of structures that a ligand has within a (default) cluster radius of 9 Å RMSD, we select the largest cluster and rank its cluster center as the top prediction of the target complex. The members of this cluster are removed from the matrix, and we select the next largest cluster and rank its center as number two, and so on. After clustering with this hierarchical approach, the ranked complexes are subjected to a straightforward (300 step and fixed backbone) minimization of the van der Waals energy using the CHARMM potential to remove potential side chain clashes.

### 2.5. Refining ClusPro results using AlphaFold2

The top 10 docked structures produced by ClusPro were provided to AlphaFold2 as templates for structure prediction. The MSA for each case was generated in the same way as described earlier (see Methods 2.2). The parameters for model 1 from the pTM parameter set was used to generate one model for each template as it gave us the best results out of the five model parameters (data not shown). This resulted in 10 refined models for each target which were ranked based on their interface PAE scores. We then also added the five models generated by AlphaFold2 in the first step of this protocol. Finally, the 15 resulting models were ranked based on their average interface PAE scores.

### 2.5. Assessing the accuracy of docked structures

We used the DockQ program to assess the interface quality of the generated models [17]. The interface of interest for each target was the one between the receptor and ligand as specified in section 2.3. DockQ outputs a score between 0 and 1 to indicate interface quality. A score < 0.23 indicates an incorrect interface, 0.23 ≤ score < 0.49 indicates an acceptable interface, 0.49 ≤ score < 0.80 indicates a medium quality interface, and score ≥ 0.80 indicates a high quality interface. A target was considered successful for a method if the interface for one of the models produced by the method was of acceptable quality or better.

## 3. Results and discussion

### 3.1. AlphaFold2 refinement improves predictions

We used the sequence information from the PDB for the 204 targets in Benchmark 1 to build the complexes using AlphaFold2. As described, the model with the lowest average interface PAE was split into two groups of residues based on the interface of interest, and docked using ClusPro. A direct comparison of the results by AlphaFold2 and ClusPro show that AlphaFold2 performs better for top 1 and top 5 predictions, both in terms of the number of targets with acceptable or better quality models and in terms of the quality of the interfaces defined by the DockQ score (**Figure 1**). However, considering the top 10 models from ClusPro increases the number of successful targets to 98, which is higher than the 88 successful targets for the 5 models from AlphaFold2. The higher success rate comes at the cost of interface quality as ClusPro provides far fewer targets with high quality interfaces.

**Figure 1:**
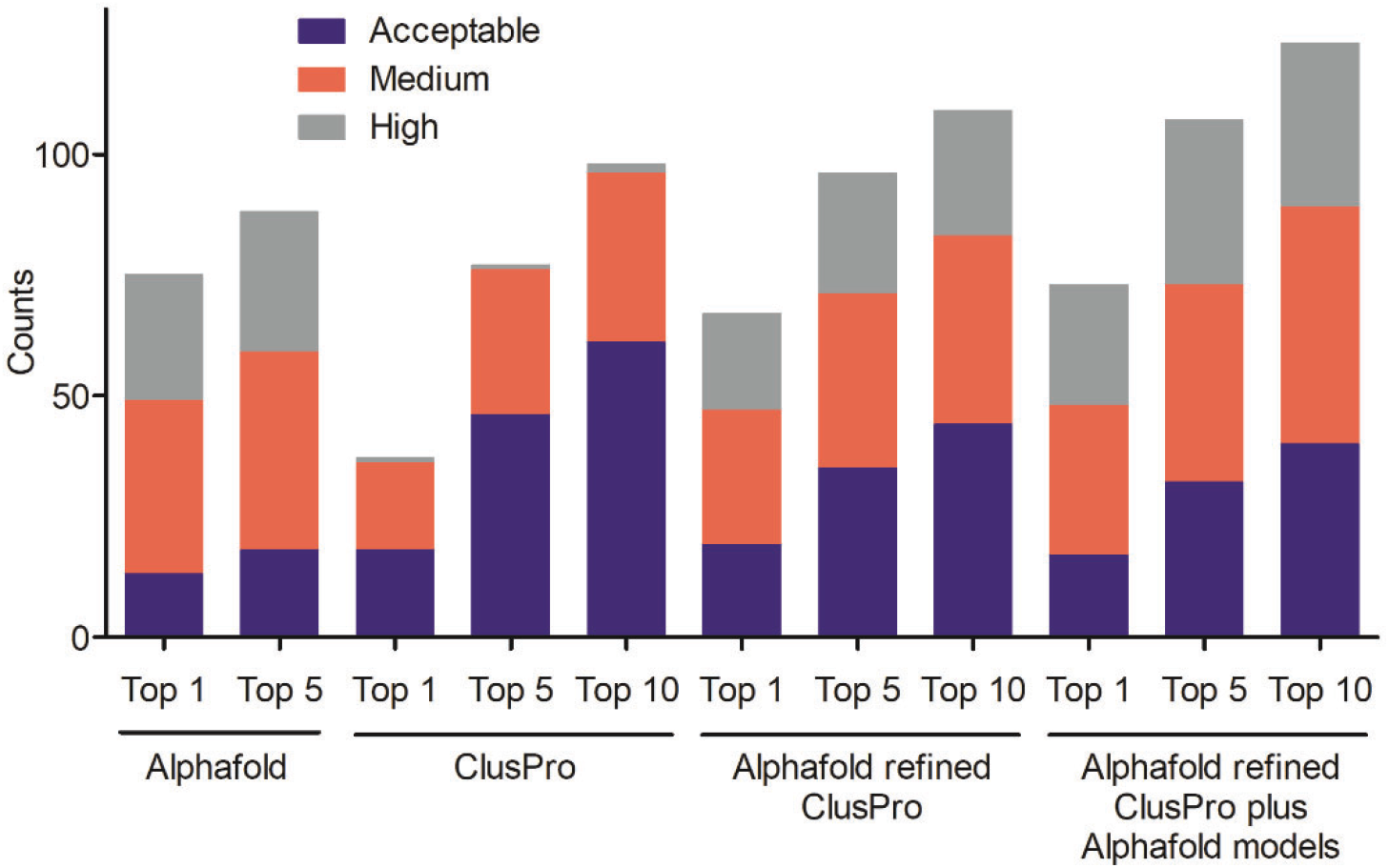
Performance on Benchmark 1. The number of successful targets and the quality of best models at the different steps of the protocol.

The 10 models generated by ClusPro were then refined using AlphaFold2 and ranked based on their average interface PAEs. As shown in **Figure 1**, this step improves the results considerably. AlphaFold2 on its own still generates good models as the top 1 prediction for more targets than the AlphaFold2 refinement of the ClusPro models. However, when considering the top 5 predictions, the AlphaFold2 refinement of the 10 ClusPro models is successful for 96 targets, compared to the 88 targets obtained by AlphaFold2 without ClusPro. Moreover, considering all 10 AlphaFold2 refined ClusPro models increases the number of successful targets to 109. For some targets where AlphaFold2 fails by itself and ClusPro produces an acceptable quality model, AlphaFold2 refinement is then able to improve the ClusPro interface to a high quality prediction (**Table S1**). However, it is also apparent that there is a loss of interface quality for some targets where AlphaFold2 by itself did better than ClusPro followed by AlphaFold2 refinement. In fact, the AlphaFold2 refinement of the 10 ClusPro models produces high quality predictions for fewer targets than when considering the top five Alphafold2 models without any further calculation (**Table S1**). To address this loss, we added the original five AlphaFold2 models to the 10 AlphaFold2 refined ClusPro models, and ranked the 15 models based on the average PAE interface scores. This improved the number of successful targets while also preserving, or even improving, the quality of the interfaces of the top models. This strategy, i.e., selecting 10 models with the lowest PAE values among the 15 models (10 AlphaFold2 refined ClusPro models plus the five original AlphaFold2 models) produced 123 successful targets, with an average DockQ score of 0.415. This was a 37.6% increase in the number of successful targets compared to ClusPro when considering the top 10 models, and almost 40% (39.8%) compared to the original AlphaFold2 runs. Thus, the combined protocol produces much higher success rates that either ClusPro or AlphaFold2 on its own. The refinement of ClusPro generated models by AlphaFold2 was particularly effective and improved the average DockQ by 58.3%. We note that the ClusPro performance discussed here was based on docking the monomer models generated by AlphaFold2, thus starting the calculations from sequences rather than structures, which is another strength of the combined protocol.

### 3.2. Refinement improvement persists for new targets

As stated, the heterodimer targets in Benchmark 2 were released in the PDB after May 2018 and did not have homologous templates. Similar to Benchmark 1, the sequences were used to derive a complex structure using Alphafold2. These were then split into subunits, docked using ClusPro, and refined by AlphaFold2. The results for these complexes differ from the results for Benchmark 1 in two significant ways. First, when considering the top 5 models, AlphaFold2 produced good predictions for fewer targets than ClusPro (**Figure 2**), although it did result in more high quality interfaces. The lower interface quality for the ClusPro results is not too surprising given that it is a rigid body docking method that does not take into account the conformational changes upon binding. It is likely that success for fewer targets by AlphaFold2 is due to the fact that these targets were specifically selected for their lack of appropriate templates in the PDB. Thus, AlphaFold2 had no or very few structures similar to the targets in the training data, reducing the quality of predictions. The second difference from the results for Benchmark 1 is that the number of successful targets did not improve when going from the ClusPro models to the AlphaFold2 refined ClusPro models, and further to the mixed version of AlphaFold2 and refined ClusPro models. However, similar to the results for Benchmark 1, interface quality improved considerably with refinement (**Figure 3**). The average DockQ score highlights this improvement as it increases from 0.422 for the top 10 ClusPro models to 0.561 for the AlphaFold2 refined ClusPro models (**Table S2**).

**Figure 2:**
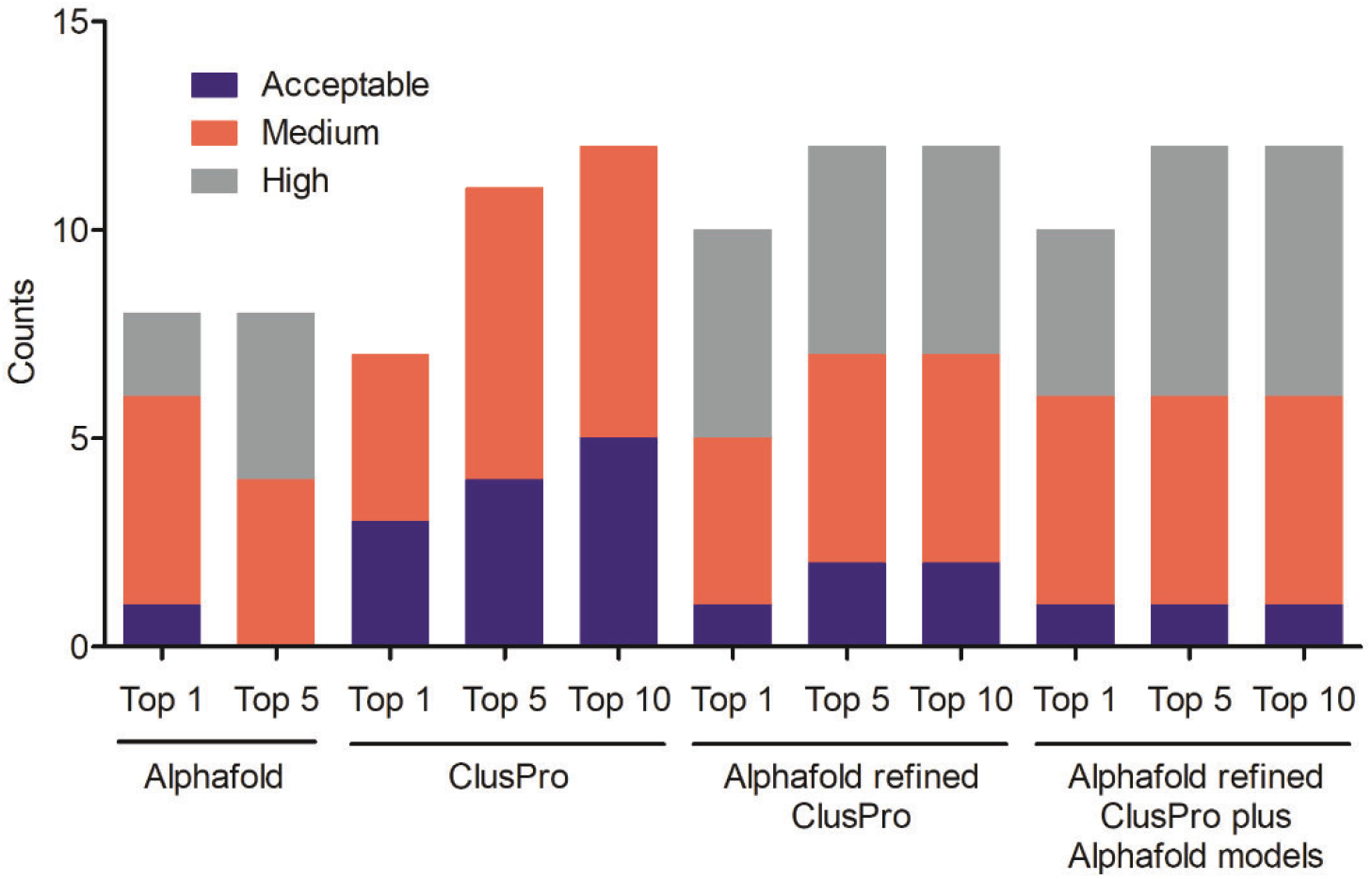
Performance on Benchmark 2. The number of successful targets and the quality of best models at the different steps of the protocol.

**Figure 3:**
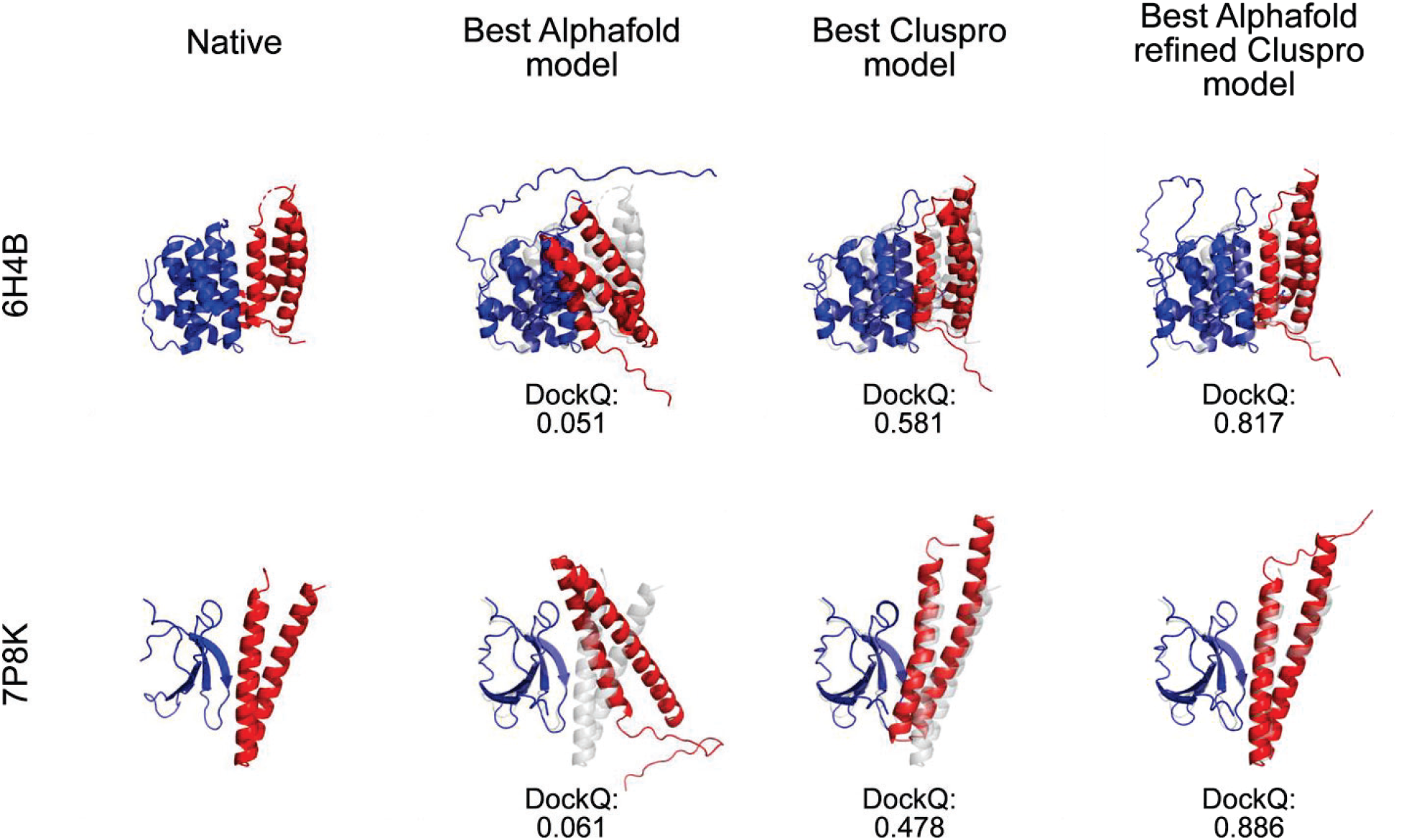
Comparison of the best model from each method to the native structure for targets 6H4B and 7P8K.

## Conclusions

We tested the ability of the neural network based protein structure prediction program AlphaFold2 of predicting the structures of protein-protein complexes. The method, established in earlier publications, is simply running the program on the two sequences separated by a gap [7]. Application to 204 complexes from the “gold standard” protein-protein docking benchmark [12] confirmed that this method yields a considerable success rate when considering the top five AlphaFold2 predictions. This performance is similar to the one observed for other protein docking programs [15]. As expected, even some of the poorly predicted complexes had excellent models of the protein monomers. The monomers were docked by the physics based traditional docking server ClusPro [11], and the resulting models were used as templates in a repeated prediction of the complexes by AlphaFold2. The combination of the two approaches substantially improved the overall performance. The performance on the protein docking benchmark (Benchmark 1 in this paper) may be somewhat biased since these proteins were part of the structures used when training the AlphaFold2 program. The possibility of such bias was eliminated by performing the same type of calculations for heterodimer complexes deposited in the PDB after May 2018, the cut-off date of training AlphaFold2, and that did not have any templates. Performance was similar to that seen for the other benchmark. Thus, we conclude that the combined protocol produces better results than either AlphaFold2 or ClusPro on its own. It is also important that the calculations can be performed without knowing the structures of the proteins to be docked, and hence we believe that the algorithm proposed here will be very useful in applications.

## Acknowledgements

This investigation was supported by grants DBI 1759277 and AF 1645512 from the National Science Foundation, and R35GM118078, R21GM127952, and RM1135136 from the National Institute of General Medical Sciences.

## Supporting Information

**Table S1:**
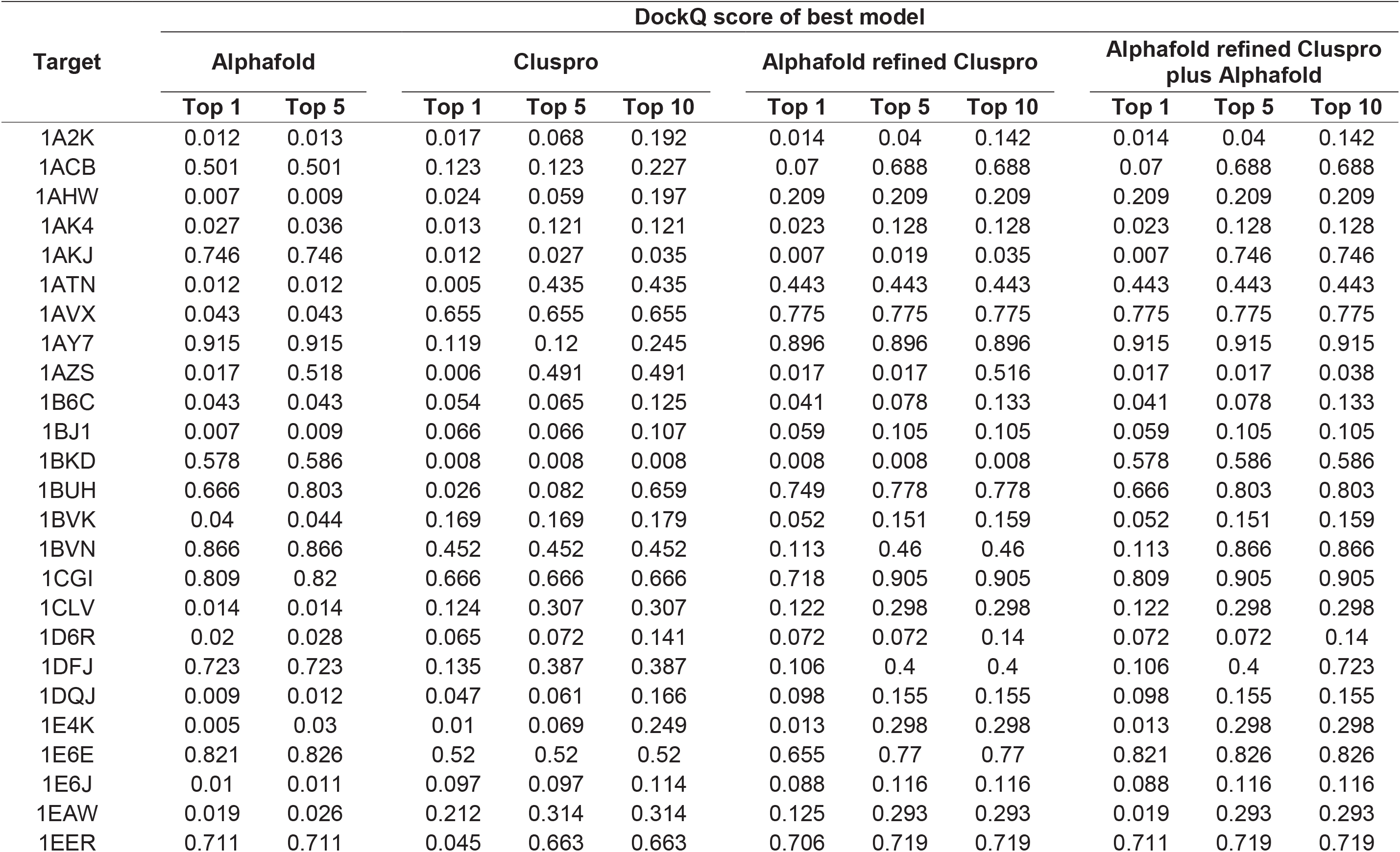
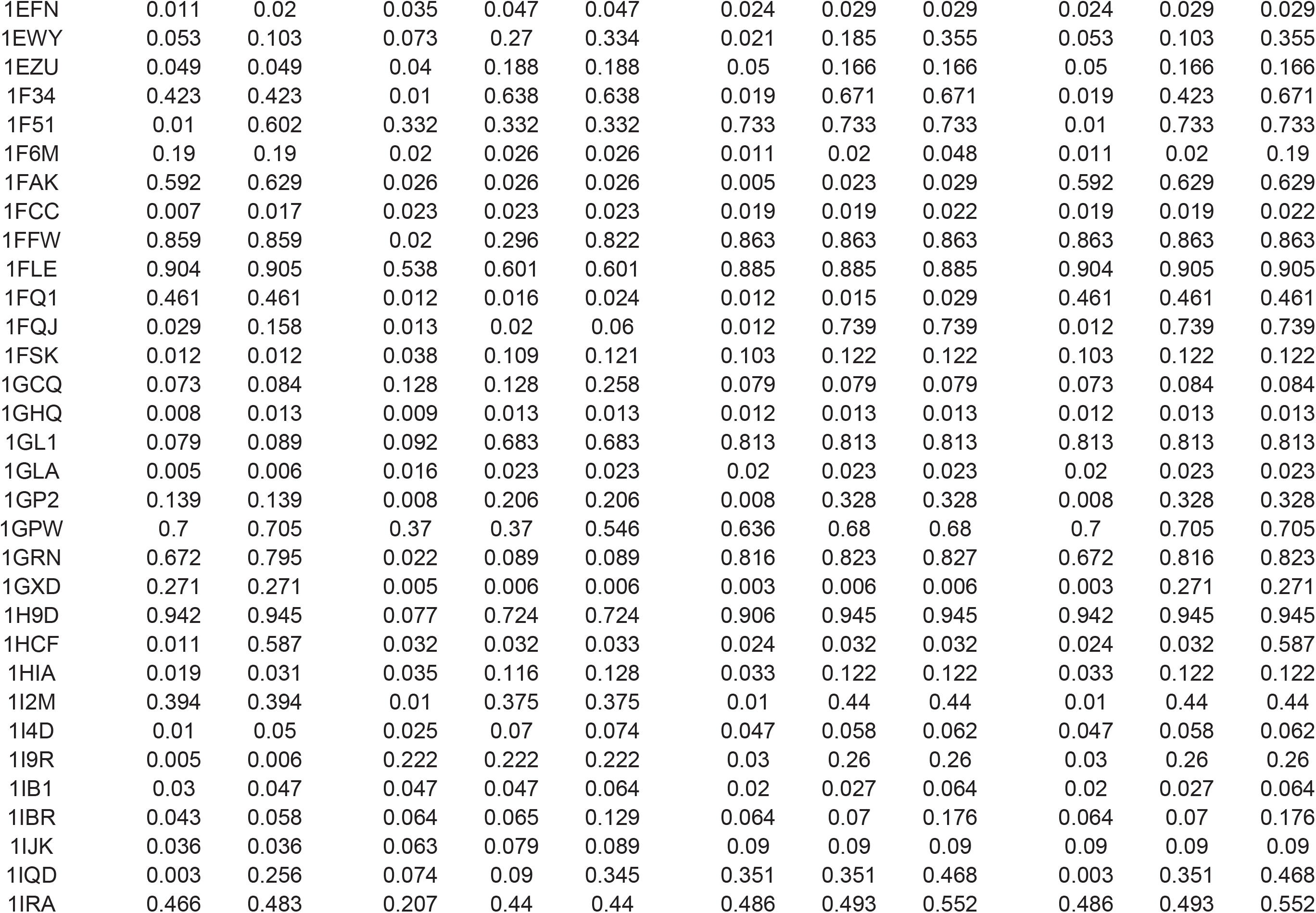
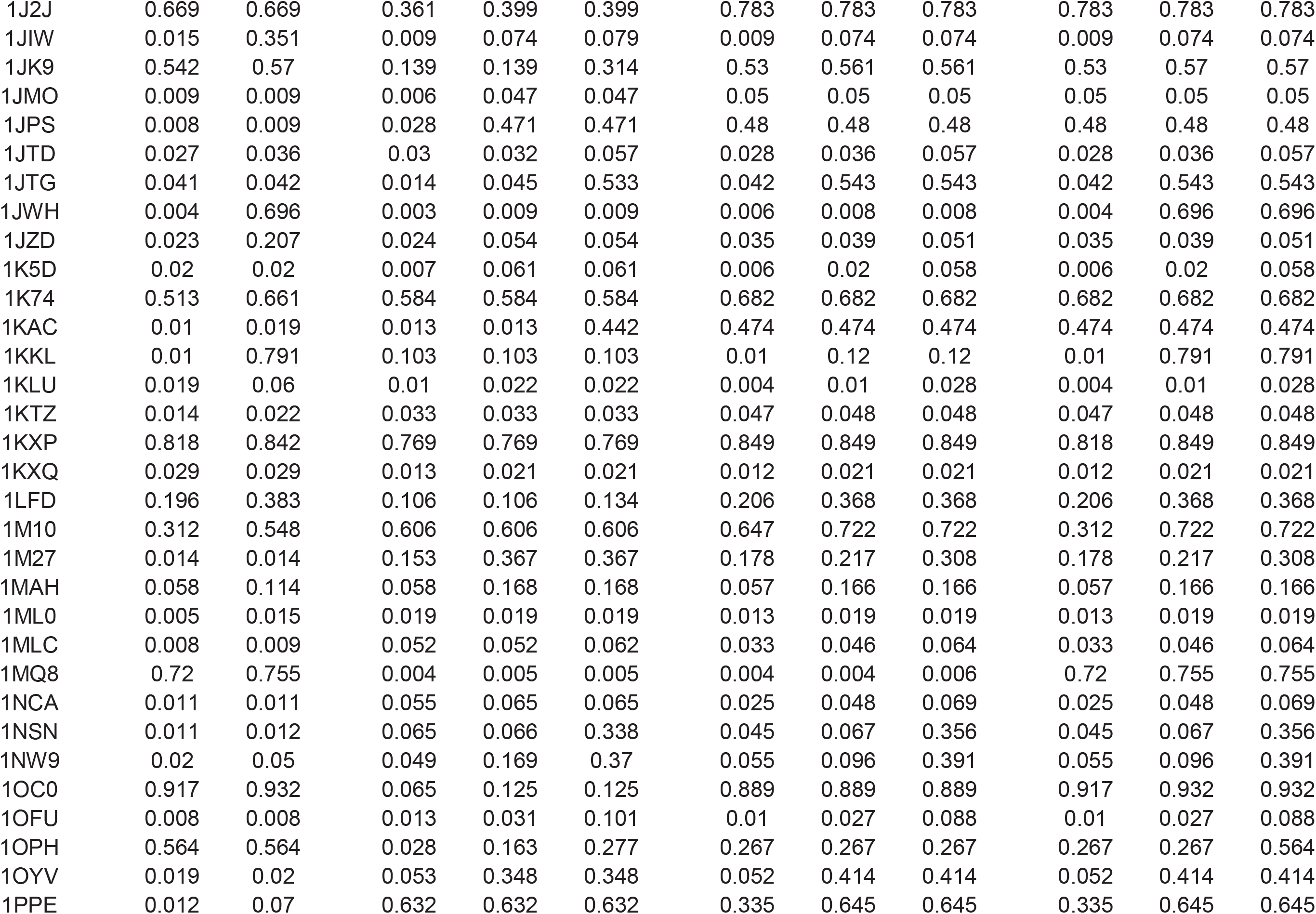
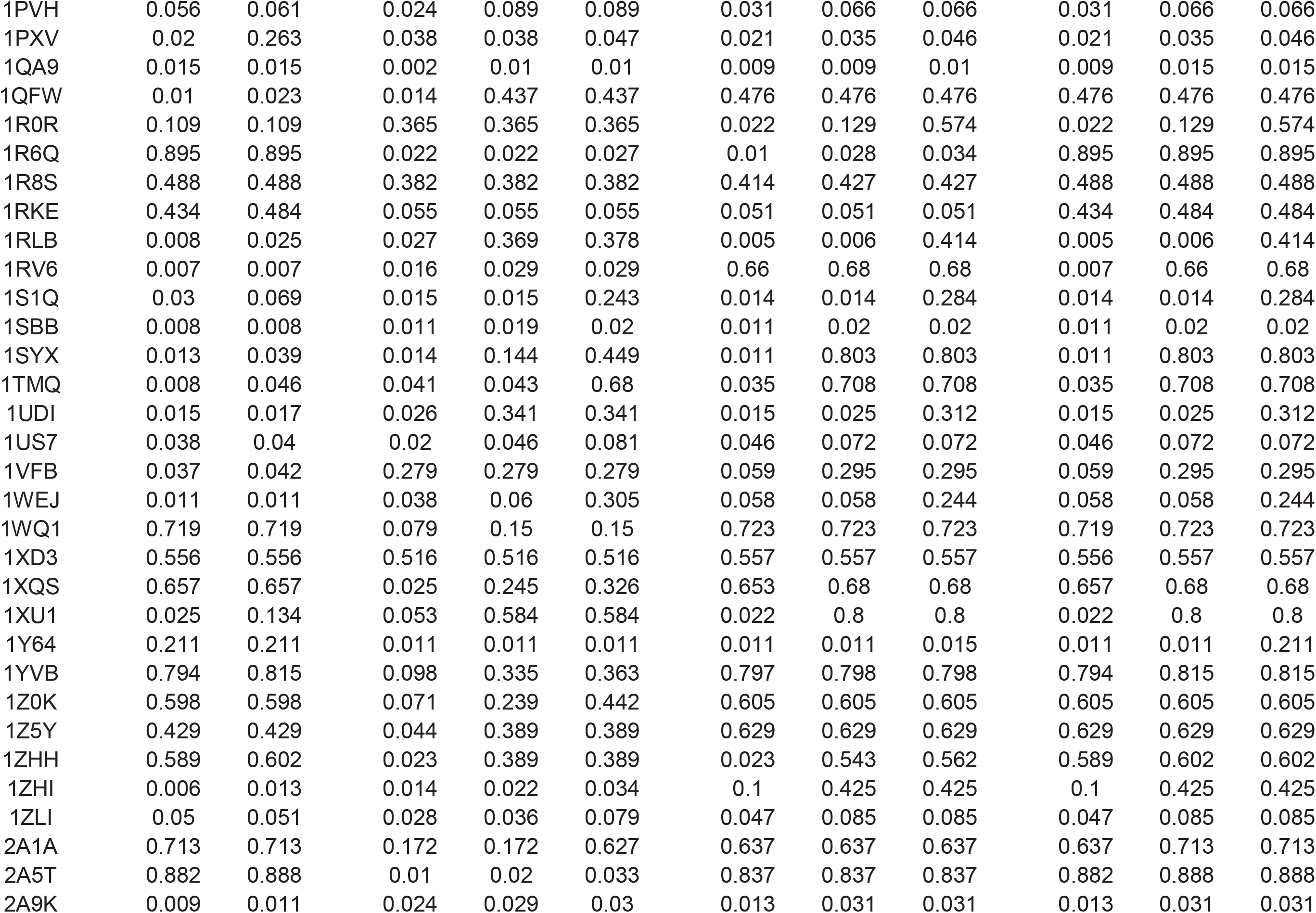
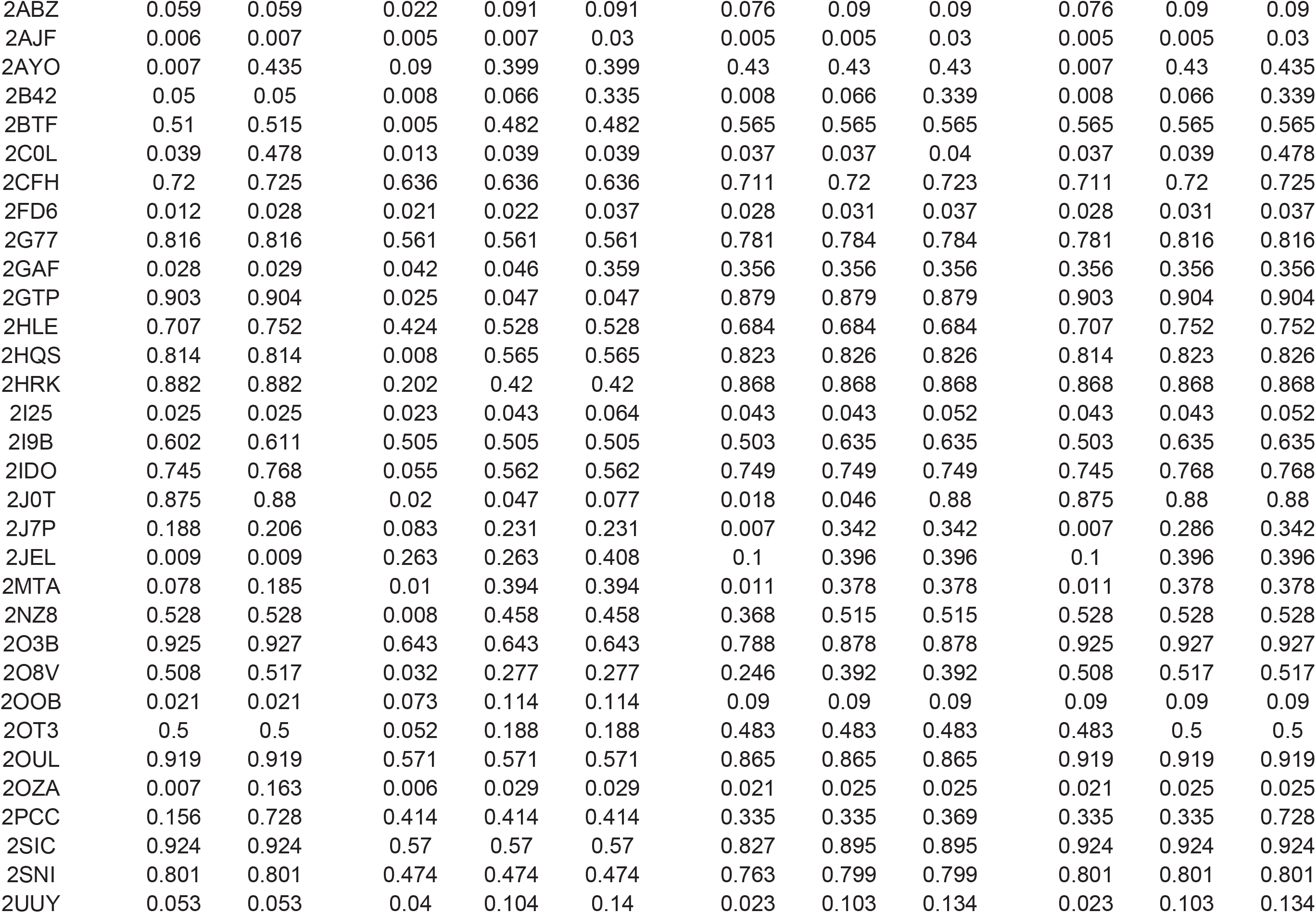
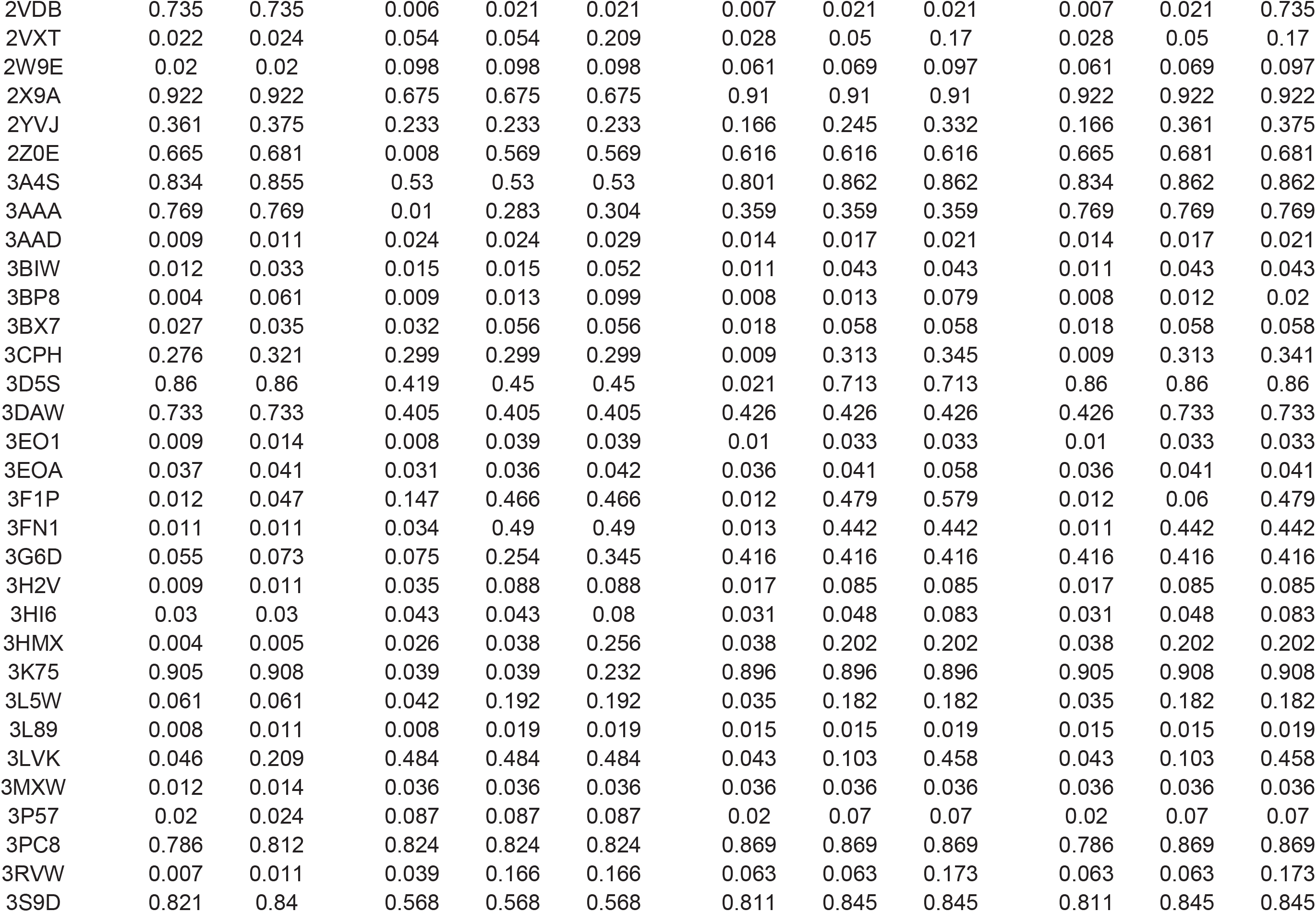
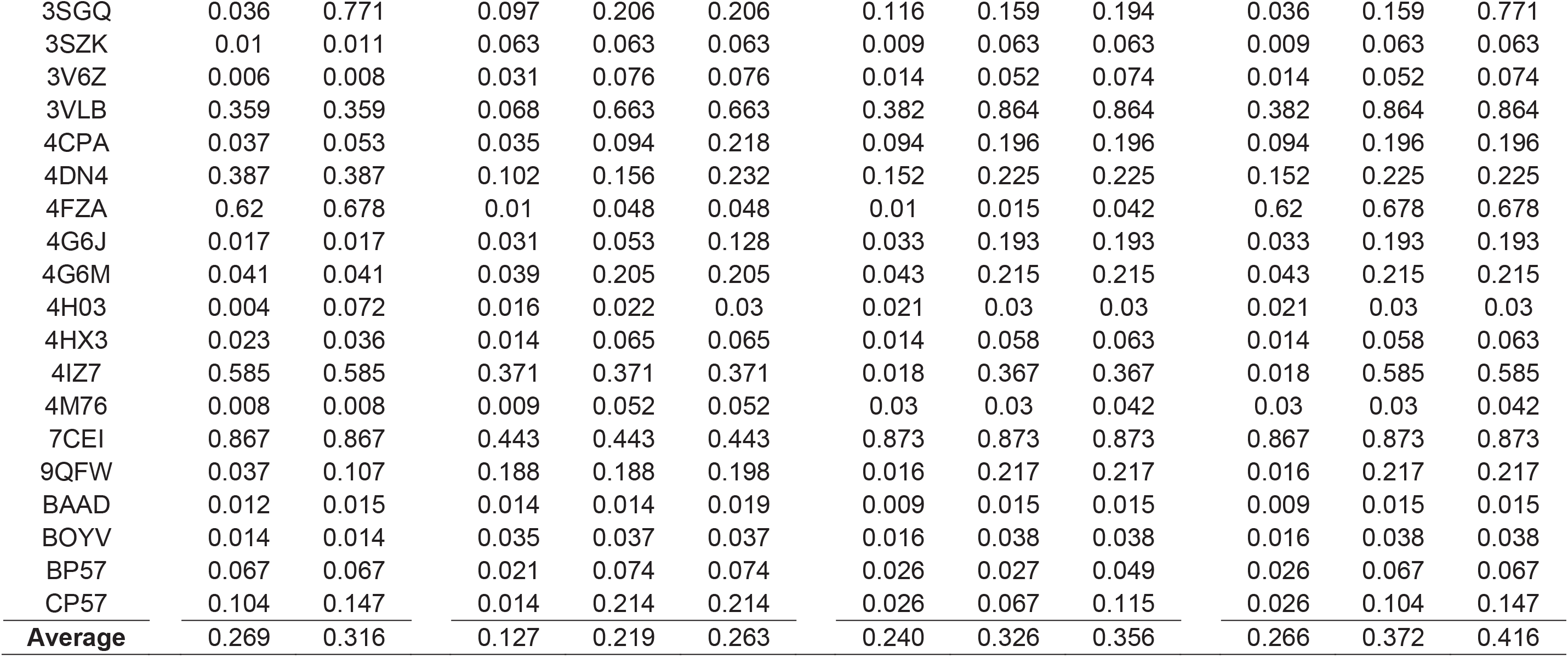
DockQ score of the best model for the targets in Benchmark 1.

| Target | DockQ score of best model |  |  |  |  |  |  |  |  |  |  |
| --- | --- | --- | --- | --- | --- | --- | --- | --- | --- | --- | --- |
|  | Alphafold |  | Cluspro |  |  | Alphafold refined Cluspro |  |  | Alphafold refined Cluspro plus Alphafold |  |  |
|  | Top 1 | Top 5 | Top 1 | Top 5 | Top 10 | Top 1 | Top 5 | Top 10 | Top 1 | Top 5 | Top 10 |
| 1A2K | 0.012 | 0.013 | 0.017 | 0.068 | 0.192 | 0.014 | 0.04 | 0.142 | 0.014 | 0.04 | 0.142 |
| 1ACB | 0.501 | 0.501 | 0.123 | 0.123 | 0.227 | 0.07 | 0.688 | 0.688 | 0.07 | 0.688 | 0.688 |
| 1AHW | 0.007 | 0.009 | 0.024 | 0.059 | 0.197 | 0.209 | 0.209 | 0.209 | 0.209 | 0.209 | 0.209 |
| 1AK4 | 0.027 | 0.036 | 0.013 | 0.121 | 0.121 | 0.023 | 0.128 | 0.128 | 0.023 | 0.128 | 0.128 |
| 1AKJ | 0.746 | 0.746 | 0.012 | 0.027 | 0.035 | 0.007 | 0.019 | 0.035 | 0.007 | 0.746 | 0.746 |
| 1ATN | 0.012 | 0.012 | 0.005 | 0.435 | 0.435 | 0.443 | 0.443 | 0.443 | 0.443 | 0.443 | 0.443 |
| 1AVX | 0.043 | 0.043 | 0.655 | 0.655 | 0.655 | 0.775 | 0.775 | 0.775 | 0.775 | 0.775 | 0.775 |
| 1AY7 | 0.915 | 0.915 | 0.119 | 0.12 | 0.245 | 0.896 | 0.896 | 0.896 | 0.915 | 0.915 | 0.915 |
| 1AZS | 0.017 | 0.518 | 0.006 | 0.491 | 0.491 | 0.017 | 0.017 | 0.516 | 0.017 | 0.017 | 0.038 |
| 1B6C | 0.043 | 0.043 | 0.054 | 0.065 | 0.125 | 0.041 | 0.078 | 0.133 | 0.041 | 0.078 | 0.133 |
| 1BJ1 | 0.007 | 0.009 | 0.066 | 0.066 | 0.107 | 0.059 | 0.105 | 0.105 | 0.059 | 0.105 | 0.105 |
| 1BKD | 0.578 | 0.586 | 0.008 | 0.008 | 0.008 | 0.008 | 0.008 | 0.008 | 0.578 | 0.586 | 0.586 |
| 1BUH | 0.666 | 0.803 | 0.026 | 0.082 | 0.659 | 0.749 | 0.778 | 0.778 | 0.666 | 0.803 | 0.803 |
| 1BVK | 0.04 | 0.044 | 0.169 | 0.169 | 0.179 | 0.052 | 0.151 | 0.159 | 0.052 | 0.151 | 0.159 |
| 1BVN | 0.866 | 0.866 | 0.452 | 0.452 | 0.452 | 0.113 | 0.46 | 0.46 | 0.113 | 0.866 | 0.866 |
| 1CGI | 0.809 | 0.82 | 0.666 | 0.666 | 0.666 | 0.718 | 0.905 | 0.905 | 0.809 | 0.905 | 0.905 |
| 1CLV | 0.014 | 0.014 | 0.124 | 0.307 | 0.307 | 0.122 | 0.298 | 0.298 | 0.122 | 0.298 | 0.298 |
| 1D6R | 0.02 | 0.028 | 0.065 | 0.072 | 0.141 | 0.072 | 0.072 | 0.14 | 0.072 | 0.072 | 0.14 |
| 1DFJ | 0.723 | 0.723 | 0.135 | 0.387 | 0.387 | 0.106 | 0.4 | 0.4 | 0.106 | 0.4 | 0.723 |
| 1DQJ | 0.009 | 0.012 | 0.047 | 0.061 | 0.166 | 0.098 | 0.155 | 0.155 | 0.098 | 0.155 | 0.155 |
| 1E4K | 0.005 | 0.03 | 0.01 | 0.069 | 0.249 | 0.013 | 0.298 | 0.298 | 0.013 | 0.298 | 0.298 |
| 1E6E | 0.821 | 0.826 | 0.52 | 0.52 | 0.52 | 0.655 | 0.77 | 0.77 | 0.821 | 0.826 | 0.826 |
| 1E6J | 0.01 | 0.011 | 0.097 | 0.097 | 0.114 | 0.088 | 0.116 | 0.116 | 0.088 | 0.116 | 0.116 |
| 1EAW | 0.019 | 0.026 | 0.212 | 0.314 | 0.314 | 0.125 | 0.293 | 0.293 | 0.019 | 0.293 | 0.293 |
| 1EER | 0.711 | 0.711 | 0.045 | 0.663 | 0.663 | 0.706 | 0.719 | 0.719 | 0.711 | 0.719 | 0.719 |
| 1EFN | 0.011 | 0.02 | 0.035 | 0.047 | 0.047 | 0.024 | 0.029 | 0.029 | 0.024 | 0.029 | 0.029 |
| 1EWY | 0.053 | 0.103 | 0.073 | 0.27 | 0.334 | 0.021 | 0.185 | 0.355 | 0.053 | 0.103 | 0.355 |
| 1EZU | 0.049 | 0.049 | 0.04 | 0.188 | 0.188 | 0.05 | 0.166 | 0.166 | 0.05 | 0.166 | 0.166 |
| 1F34 | 0.423 | 0.423 | 0.01 | 0.638 | 0.638 | 0.019 | 0.671 | 0.671 | 0.019 | 0.423 | 0.671 |
| 1F51 | 0.01 | 0.602 | 0.332 | 0.332 | 0.332 | 0.733 | 0.733 | 0.733 | 0.01 | 0.733 | 0.733 |
| 1F6M | 0.19 | 0.19 | 0.02 | 0.026 | 0.026 | 0.011 | 0.02 | 0.048 | 0.011 | 0.02 | 0.19 |
| 1FAK | 0.592 | 0.629 | 0.026 | 0.026 | 0.026 | 0.005 | 0.023 | 0.029 | 0.592 | 0.629 | 0.629 |
| 1FCC | 0.007 | 0.017 | 0.023 | 0.023 | 0.023 | 0.019 | 0.019 | 0.022 | 0.019 | 0.019 | 0.022 |
| 1FFW | 0.859 | 0.859 | 0.02 | 0.296 | 0.822 | 0.863 | 0.863 | 0.863 | 0.863 | 0.863 | 0.863 |
| 1FLE | 0.904 | 0.905 | 0.538 | 0.601 | 0.601 | 0.885 | 0.885 | 0.885 | 0.904 | 0.905 | 0.905 |
| 1FQ1 | 0.461 | 0.461 | 0.012 | 0.016 | 0.024 | 0.012 | 0.015 | 0.029 | 0.461 | 0.461 | 0.461 |
| 1FQJ | 0.029 | 0.158 | 0.013 | 0.02 | 0.06 | 0.012 | 0.739 | 0.739 | 0.012 | 0.739 | 0.739 |
| 1FSK | 0.012 | 0.012 | 0.038 | 0.109 | 0.121 | 0.103 | 0.122 | 0.122 | 0.103 | 0.122 | 0.122 |
| 1GCQ | 0.073 | 0.084 | 0.128 | 0.128 | 0.258 | 0.079 | 0.079 | 0.079 | 0.073 | 0.084 | 0.084 |
| 1GHQ | 0.008 | 0.013 | 0.009 | 0.013 | 0.013 | 0.012 | 0.013 | 0.013 | 0.012 | 0.013 | 0.013 |
| 1GL1 | 0.079 | 0.089 | 0.092 | 0.683 | 0.683 | 0.813 | 0.813 | 0.813 | 0.813 | 0.813 | 0.813 |
| 1GLA | 0.005 | 0.006 | 0.016 | 0.023 | 0.023 | 0.02 | 0.023 | 0.023 | 0.02 | 0.023 | 0.023 |
| 1GP2 | 0.139 | 0.139 | 0.008 | 0.206 | 0.206 | 0.008 | 0.328 | 0.328 | 0.008 | 0.328 | 0.328 |
| 1GPW | 0.7 | 0.705 | 0.37 | 0.37 | 0.546 | 0.636 | 0.68 | 0.68 | 0.7 | 0.705 | 0.705 |
| 1GRN | 0.672 | 0.795 | 0.022 | 0.089 | 0.089 | 0.816 | 0.823 | 0.827 | 0.672 | 0.816 | 0.823 |
| 1GXD | 0.271 | 0.271 | 0.005 | 0.006 | 0.006 | 0.003 | 0.006 | 0.006 | 0.003 | 0.271 | 0.271 |
| 1H9D | 0.942 | 0.945 | 0.077 | 0.724 | 0.724 | 0.906 | 0.945 | 0.945 | 0.942 | 0.945 | 0.945 |
| 1HCF | 0.011 | 0.587 | 0.032 | 0.032 | 0.033 | 0.024 | 0.032 | 0.032 | 0.024 | 0.032 | 0.587 |
| 1HIA | 0.019 | 0.031 | 0.035 | 0.116 | 0.128 | 0.033 | 0.122 | 0.122 | 0.033 | 0.122 | 0.122 |
| 1I2M | 0.394 | 0.394 | 0.01 | 0.375 | 0.375 | 0.01 | 0.44 | 0.44 | 0.01 | 0.44 | 0.44 |
| 1I4D | 0.01 | 0.05 | 0.025 | 0.07 | 0.074 | 0.047 | 0.058 | 0.062 | 0.047 | 0.058 | 0.062 |
| 1I9R | 0.005 | 0.006 | 0.222 | 0.222 | 0.222 | 0.03 | 0.26 | 0.26 | 0.03 | 0.26 | 0.26 |
| 1IB1 | 0.03 | 0.047 | 0.047 | 0.047 | 0.064 | 0.02 | 0.027 | 0.064 | 0.02 | 0.027 | 0.064 |
| 1IBR | 0.043 | 0.058 | 0.064 | 0.065 | 0.129 | 0.064 | 0.07 | 0.176 | 0.064 | 0.07 | 0.176 |
| 1IJK | 0.036 | 0.036 | 0.063 | 0.079 | 0.089 | 0.09 | 0.09 | 0.09 | 0.09 | 0.09 | 0.09 |
| 1IQD | 0.003 | 0.256 | 0.074 | 0.09 | 0.345 | 0.351 | 0.351 | 0.468 | 0.003 | 0.351 | 0.468 |
| 1IRA | 0.466 | 0.483 | 0.207 | 0.44 | 0.44 | 0.486 | 0.493 | 0.552 | 0.486 | 0.493 | 0.552 |
| 1J2J | 0.669 | 0.669 | 0.361 | 0.399 | 0.399 | 0.783 | 0.783 | 0.783 | 0.783 | 0.783 | 0.783 |
| 1JIW | 0.015 | 0.351 | 0.009 | 0.074 | 0.079 | 0.009 | 0.074 | 0.074 | 0.009 | 0.074 | 0.074 |
| 1JK9 | 0.542 | 0.57 | 0.139 | 0.139 | 0.314 | 0.53 | 0.561 | 0.561 | 0.53 | 0.57 | 0.57 |
| 1JMO | 0.009 | 0.009 | 0.006 | 0.047 | 0.047 | 0.05 | 0.05 | 0.05 | 0.05 | 0.05 | 0.05 |
| 1JPS | 0.008 | 0.009 | 0.028 | 0.471 | 0.471 | 0.48 | 0.48 | 0.48 | 0.48 | 0.48 | 0.48 |
| 1JTD | 0.027 | 0.036 | 0.03 | 0.032 | 0.057 | 0.028 | 0.036 | 0.057 | 0.028 | 0.036 | 0.057 |
| 1JTG | 0.041 | 0.042 | 0.014 | 0.045 | 0.533 | 0.042 | 0.543 | 0.543 | 0.042 | 0.543 | 0.543 |
| 1JWH | 0.004 | 0.696 | 0.003 | 0.009 | 0.009 | 0.006 | 0.008 | 0.008 | 0.004 | 0.696 | 0.696 |
| 1JZD | 0.023 | 0.207 | 0.024 | 0.054 | 0.054 | 0.035 | 0.039 | 0.051 | 0.035 | 0.039 | 0.051 |
| 1K5D | 0.02 | 0.02 | 0.007 | 0.061 | 0.061 | 0.006 | 0.02 | 0.058 | 0.006 | 0.02 | 0.058 |
| 1K74 | 0.513 | 0.661 | 0.584 | 0.584 | 0.584 | 0.682 | 0.682 | 0.682 | 0.682 | 0.682 | 0.682 |
| 1KAC | 0.01 | 0.019 | 0.013 | 0.013 | 0.442 | 0.474 | 0.474 | 0.474 | 0.474 | 0.474 | 0.474 |
| 1KKL | 0.01 | 0.791 | 0.103 | 0.103 | 0.103 | 0.01 | 0.12 | 0.12 | 0.01 | 0.791 | 0.791 |
| 1KLU | 0.019 | 0.06 | 0.01 | 0.022 | 0.022 | 0.004 | 0.01 | 0.028 | 0.004 | 0.01 | 0.028 |
| 1KTZ | 0.014 | 0.022 | 0.033 | 0.033 | 0.033 | 0.047 | 0.048 | 0.048 | 0.047 | 0.048 | 0.048 |
| 1KXP | 0.818 | 0.842 | 0.769 | 0.769 | 0.769 | 0.849 | 0.849 | 0.849 | 0.818 | 0.849 | 0.849 |
| 1KXQ | 0.029 | 0.029 | 0.013 | 0.021 | 0.021 | 0.012 | 0.021 | 0.021 | 0.012 | 0.021 | 0.021 |
| 1LFD | 0.196 | 0.383 | 0.106 | 0.106 | 0.134 | 0.206 | 0.368 | 0.368 | 0.206 | 0.368 | 0.368 |
| 1M10 | 0.312 | 0.548 | 0.606 | 0.606 | 0.606 | 0.647 | 0.722 | 0.722 | 0.312 | 0.722 | 0.722 |
| 1M27 | 0.014 | 0.014 | 0.153 | 0.367 | 0.367 | 0.178 | 0.217 | 0.308 | 0.178 | 0.217 | 0.308 |
| 1MAH | 0.058 | 0.114 | 0.058 | 0.168 | 0.168 | 0.057 | 0.166 | 0.166 | 0.057 | 0.166 | 0.166 |
| 1ML0 | 0.005 | 0.015 | 0.019 | 0.019 | 0.019 | 0.013 | 0.019 | 0.019 | 0.013 | 0.019 | 0.019 |
| 1MLC | 0.008 | 0.009 | 0.052 | 0.052 | 0.062 | 0.033 | 0.046 | 0.064 | 0.033 | 0.046 | 0.064 |
| 1MQ8 | 0.72 | 0.755 | 0.004 | 0.005 | 0.005 | 0.004 | 0.004 | 0.006 | 0.72 | 0.755 | 0.755 |
| 1NCA | 0.011 | 0.011 | 0.055 | 0.065 | 0.065 | 0.025 | 0.048 | 0.069 | 0.025 | 0.048 | 0.069 |
| 1NSN | 0.011 | 0.012 | 0.065 | 0.066 | 0.338 | 0.045 | 0.067 | 0.356 | 0.045 | 0.067 | 0.356 |
| 1NW9 | 0.02 | 0.05 | 0.049 | 0.169 | 0.37 | 0.055 | 0.096 | 0.391 | 0.055 | 0.096 | 0.391 |
| 1OC0 | 0.917 | 0.932 | 0.065 | 0.125 | 0.125 | 0.889 | 0.889 | 0.889 | 0.917 | 0.932 | 0.932 |
| 1OFU | 0.008 | 0.008 | 0.013 | 0.031 | 0.101 | 0.01 | 0.027 | 0.088 | 0.01 | 0.027 | 0.088 |
| 1OPH | 0.564 | 0.564 | 0.028 | 0.163 | 0.277 | 0.267 | 0.267 | 0.267 | 0.267 | 0.267 | 0.564 |
| 1OYV | 0.019 | 0.02 | 0.053 | 0.348 | 0.348 | 0.052 | 0.414 | 0.414 | 0.052 | 0.414 | 0.414 |
| 1PPE | 0.012 | 0.07 | 0.632 | 0.632 | 0.632 | 0.335 | 0.645 | 0.645 | 0.335 | 0.645 | 0.645 |
| 1PVH | 0.056 | 0.061 | 0.024 | 0.089 | 0.089 | 0.031 | 0.066 | 0.066 | 0.031 | 0.066 | 0.066 |
| 1PXV | 0.02 | 0.263 | 0.038 | 0.038 | 0.047 | 0.021 | 0.035 | 0.046 | 0.021 | 0.035 | 0.046 |
| 1QA9 | 0.015 | 0.015 | 0.002 | 0.01 | 0.01 | 0.009 | 0.009 | 0.01 | 0.009 | 0.015 | 0.015 |
| 1QFW | 0.01 | 0.023 | 0.014 | 0.437 | 0.437 | 0.476 | 0.476 | 0.476 | 0.476 | 0.476 | 0.476 |
| 1R0R | 0.109 | 0.109 | 0.365 | 0.365 | 0.365 | 0.022 | 0.129 | 0.574 | 0.022 | 0.129 | 0.574 |
| 1R6Q | 0.895 | 0.895 | 0.022 | 0.022 | 0.027 | 0.01 | 0.028 | 0.034 | 0.895 | 0.895 | 0.895 |
| 1R8S | 0.488 | 0.488 | 0.382 | 0.382 | 0.382 | 0.414 | 0.427 | 0.427 | 0.488 | 0.488 | 0.488 |
| 1RKE | 0.434 | 0.484 | 0.055 | 0.055 | 0.055 | 0.051 | 0.051 | 0.051 | 0.434 | 0.484 | 0.484 |
| 1RLB | 0.008 | 0.025 | 0.027 | 0.369 | 0.378 | 0.005 | 0.006 | 0.414 | 0.005 | 0.006 | 0.414 |
| 1RV6 | 0.007 | 0.007 | 0.016 | 0.029 | 0.029 | 0.66 | 0.68 | 0.68 | 0.007 | 0.66 | 0.68 |
| 1S1Q | 0.03 | 0.069 | 0.015 | 0.015 | 0.243 | 0.014 | 0.014 | 0.284 | 0.014 | 0.014 | 0.284 |
| 1SBB | 0.008 | 0.008 | 0.011 | 0.019 | 0.02 | 0.011 | 0.02 | 0.02 | 0.011 | 0.02 | 0.02 |
| 1SYX | 0.013 | 0.039 | 0.014 | 0.144 | 0.449 | 0.011 | 0.803 | 0.803 | 0.011 | 0.803 | 0.803 |
| 1TMQ | 0.008 | 0.046 | 0.041 | 0.043 | 0.68 | 0.035 | 0.708 | 0.708 | 0.035 | 0.708 | 0.708 |
| 1UDI | 0.015 | 0.017 | 0.026 | 0.341 | 0.341 | 0.015 | 0.025 | 0.312 | 0.015 | 0.025 | 0.312 |
| 1US7 | 0.038 | 0.04 | 0.02 | 0.046 | 0.081 | 0.046 | 0.072 | 0.072 | 0.046 | 0.072 | 0.072 |
| 1VFB | 0.037 | 0.042 | 0.279 | 0.279 | 0.279 | 0.059 | 0.295 | 0.295 | 0.059 | 0.295 | 0.295 |
| 1WEJ | 0.011 | 0.011 | 0.038 | 0.06 | 0.305 | 0.058 | 0.058 | 0.244 | 0.058 | 0.058 | 0.244 |
| 1WQ1 | 0.719 | 0.719 | 0.079 | 0.15 | 0.15 | 0.723 | 0.723 | 0.723 | 0.719 | 0.723 | 0.723 |
| 1XD3 | 0.556 | 0.556 | 0.516 | 0.516 | 0.516 | 0.557 | 0.557 | 0.557 | 0.556 | 0.557 | 0.557 |
| 1XQS | 0.657 | 0.657 | 0.025 | 0.245 | 0.326 | 0.653 | 0.68 | 0.68 | 0.657 | 0.68 | 0.68 |
| 1XU1 | 0.025 | 0.134 | 0.053 | 0.584 | 0.584 | 0.022 | 0.8 | 0.8 | 0.022 | 0.8 | 0.8 |
| 1Y64 | 0.211 | 0.211 | 0.011 | 0.011 | 0.011 | 0.011 | 0.011 | 0.015 | 0.011 | 0.011 | 0.211 |
| 1YVB | 0.794 | 0.815 | 0.098 | 0.335 | 0.363 | 0.797 | 0.798 | 0.798 | 0.794 | 0.815 | 0.815 |
| 1Z0K | 0.598 | 0.598 | 0.071 | 0.239 | 0.442 | 0.605 | 0.605 | 0.605 | 0.605 | 0.605 | 0.605 |
| 1Z5Y | 0.429 | 0.429 | 0.044 | 0.389 | 0.389 | 0.629 | 0.629 | 0.629 | 0.629 | 0.629 | 0.629 |
| 1ZHH | 0.589 | 0.602 | 0.023 | 0.389 | 0.389 | 0.023 | 0.543 | 0.562 | 0.589 | 0.602 | 0.602 |
| 1ZHI | 0.006 | 0.013 | 0.014 | 0.022 | 0.034 | 0.1 | 0.425 | 0.425 | 0.1 | 0.425 | 0.425 |
| 1ZLI | 0.05 | 0.051 | 0.028 | 0.036 | 0.079 | 0.047 | 0.085 | 0.085 | 0.047 | 0.085 | 0.085 |
| 2A1A | 0.713 | 0.713 | 0.172 | 0.172 | 0.627 | 0.637 | 0.637 | 0.637 | 0.637 | 0.713 | 0.713 |
| 2A5T | 0.882 | 0.888 | 0.01 | 0.02 | 0.033 | 0.837 | 0.837 | 0.837 | 0.882 | 0.888 | 0.888 |
| 2A9K | 0.009 | 0.011 | 0.024 | 0.029 | 0.03 | 0.013 | 0.031 | 0.031 | 0.013 | 0.031 | 0.031 |
| 2ABZ | 0.059 | 0.059 | 0.022 | 0.091 | 0.091 | 0.076 | 0.09 | 0.09 | 0.076 | 0.09 | 0.09 |
| 2AJF | 0.006 | 0.007 | 0.005 | 0.007 | 0.03 | 0.005 | 0.005 | 0.03 | 0.005 | 0.005 | 0.03 |
| 2AYO | 0.007 | 0.435 | 0.09 | 0.399 | 0.399 | 0.43 | 0.43 | 0.43 | 0.007 | 0.43 | 0.435 |
| 2B42 | 0.05 | 0.05 | 0.008 | 0.066 | 0.335 | 0.008 | 0.066 | 0.339 | 0.008 | 0.066 | 0.339 |
| 2BTF | 0.51 | 0.515 | 0.005 | 0.482 | 0.482 | 0.565 | 0.565 | 0.565 | 0.565 | 0.565 | 0.565 |
| 2C0L | 0.039 | 0.478 | 0.013 | 0.039 | 0.039 | 0.037 | 0.037 | 0.04 | 0.037 | 0.039 | 0.478 |
| 2CFH | 0.72 | 0.725 | 0.636 | 0.636 | 0.636 | 0.711 | 0.72 | 0.723 | 0.711 | 0.72 | 0.725 |
| 2FD6 | 0.012 | 0.028 | 0.021 | 0.022 | 0.037 | 0.028 | 0.031 | 0.037 | 0.028 | 0.031 | 0.037 |
| 2G77 | 0.816 | 0.816 | 0.561 | 0.561 | 0.561 | 0.781 | 0.784 | 0.784 | 0.781 | 0.816 | 0.816 |
| 2GAF | 0.028 | 0.029 | 0.042 | 0.046 | 0.359 | 0.356 | 0.356 | 0.356 | 0.356 | 0.356 | 0.356 |
| 2GTP | 0.903 | 0.904 | 0.025 | 0.047 | 0.047 | 0.879 | 0.879 | 0.879 | 0.903 | 0.904 | 0.904 |
| 2HLE | 0.707 | 0.752 | 0.424 | 0.528 | 0.528 | 0.684 | 0.684 | 0.684 | 0.707 | 0.752 | 0.752 |
| 2HQS | 0.814 | 0.814 | 0.008 | 0.565 | 0.565 | 0.823 | 0.826 | 0.826 | 0.814 | 0.823 | 0.826 |
| 2HRK | 0.882 | 0.882 | 0.202 | 0.42 | 0.42 | 0.868 | 0.868 | 0.868 | 0.868 | 0.868 | 0.868 |
| 2I25 | 0.025 | 0.025 | 0.023 | 0.043 | 0.064 | 0.043 | 0.043 | 0.052 | 0.043 | 0.043 | 0.052 |
| 2I9B | 0.602 | 0.611 | 0.505 | 0.505 | 0.505 | 0.503 | 0.635 | 0.635 | 0.503 | 0.635 | 0.635 |
| 2IDO | 0.745 | 0.768 | 0.055 | 0.562 | 0.562 | 0.749 | 0.749 | 0.749 | 0.745 | 0.768 | 0.768 |
| 2J0T | 0.875 | 0.88 | 0.02 | 0.047 | 0.077 | 0.018 | 0.046 | 0.88 | 0.875 | 0.88 | 0.88 |
| 2J7P | 0.188 | 0.206 | 0.083 | 0.231 | 0.231 | 0.007 | 0.342 | 0.342 | 0.007 | 0.286 | 0.342 |
| 2JEL | 0.009 | 0.009 | 0.263 | 0.263 | 0.408 | 0.1 | 0.396 | 0.396 | 0.1 | 0.396 | 0.396 |
| 2MTA | 0.078 | 0.185 | 0.01 | 0.394 | 0.394 | 0.011 | 0.378 | 0.378 | 0.011 | 0.378 | 0.378 |
| 2NZ8 | 0.528 | 0.528 | 0.008 | 0.458 | 0.458 | 0.368 | 0.515 | 0.515 | 0.528 | 0.528 | 0.528 |
| 2O3B | 0.925 | 0.927 | 0.643 | 0.643 | 0.643 | 0.788 | 0.878 | 0.878 | 0.925 | 0.927 | 0.927 |
| 2O8V | 0.508 | 0.517 | 0.032 | 0.277 | 0.277 | 0.246 | 0.392 | 0.392 | 0.508 | 0.517 | 0.517 |
| 2OOB | 0.021 | 0.021 | 0.073 | 0.114 | 0.114 | 0.09 | 0.09 | 0.09 | 0.09 | 0.09 | 0.09 |
| 2OT3 | 0.5 | 0.5 | 0.052 | 0.188 | 0.188 | 0.483 | 0.483 | 0.483 | 0.483 | 0.5 | 0.5 |
| 2OUL | 0.919 | 0.919 | 0.571 | 0.571 | 0.571 | 0.865 | 0.865 | 0.865 | 0.919 | 0.919 | 0.919 |
| 2OZA | 0.007 | 0.163 | 0.006 | 0.029 | 0.029 | 0.021 | 0.025 | 0.025 | 0.021 | 0.025 | 0.025 |
| 2PCC | 0.156 | 0.728 | 0.414 | 0.414 | 0.414 | 0.335 | 0.335 | 0.369 | 0.335 | 0.335 | 0.728 |
| 2SIC | 0.924 | 0.924 | 0.57 | 0.57 | 0.57 | 0.827 | 0.895 | 0.895 | 0.924 | 0.924 | 0.924 |
| 2SNI | 0.801 | 0.801 | 0.474 | 0.474 | 0.474 | 0.763 | 0.799 | 0.799 | 0.801 | 0.801 | 0.801 |
| 2UUY | 0.053 | 0.053 | 0.04 | 0.104 | 0.14 | 0.023 | 0.103 | 0.134 | 0.023 | 0.103 | 0.134 |
| 2VDB | 0.735 | 0.735 | 0.006 | 0.021 | 0.021 | 0.007 | 0.021 | 0.021 | 0.007 | 0.021 | 0.735 |
| 2VXT | 0.022 | 0.024 | 0.054 | 0.054 | 0.209 | 0.028 | 0.05 | 0.17 | 0.028 | 0.05 | 0.17 |
| 2W9E | 0.02 | 0.02 | 0.098 | 0.098 | 0.098 | 0.061 | 0.069 | 0.097 | 0.061 | 0.069 | 0.097 |
| 2X9A | 0.922 | 0.922 | 0.675 | 0.675 | 0.675 | 0.91 | 0.91 | 0.91 | 0.922 | 0.922 | 0.922 |
| 2YVJ | 0.361 | 0.375 | 0.233 | 0.233 | 0.233 | 0.166 | 0.245 | 0.332 | 0.166 | 0.361 | 0.375 |
| 2Z0E | 0.665 | 0.681 | 0.008 | 0.569 | 0.569 | 0.616 | 0.616 | 0.616 | 0.665 | 0.681 | 0.681 |
| 3A4S | 0.834 | 0.855 | 0.53 | 0.53 | 0.53 | 0.801 | 0.862 | 0.862 | 0.834 | 0.862 | 0.862 |
| 3AAA | 0.769 | 0.769 | 0.01 | 0.283 | 0.304 | 0.359 | 0.359 | 0.359 | 0.769 | 0.769 | 0.769 |
| 3AAD | 0.009 | 0.011 | 0.024 | 0.024 | 0.029 | 0.014 | 0.017 | 0.021 | 0.014 | 0.017 | 0.021 |
| 3BIW | 0.012 | 0.033 | 0.015 | 0.015 | 0.052 | 0.011 | 0.043 | 0.043 | 0.011 | 0.043 | 0.043 |
| 3BP8 | 0.004 | 0.061 | 0.009 | 0.013 | 0.099 | 0.008 | 0.013 | 0.079 | 0.008 | 0.012 | 0.02 |
| 3BX7 | 0.027 | 0.035 | 0.032 | 0.056 | 0.056 | 0.018 | 0.058 | 0.058 | 0.018 | 0.058 | 0.058 |
| 3CPH | 0.276 | 0.321 | 0.299 | 0.299 | 0.299 | 0.009 | 0.313 | 0.345 | 0.009 | 0.313 | 0.341 |
| 3D5S | 0.86 | 0.86 | 0.419 | 0.45 | 0.45 | 0.021 | 0.713 | 0.713 | 0.86 | 0.86 | 0.86 |
| 3DAW | 0.733 | 0.733 | 0.405 | 0.405 | 0.405 | 0.426 | 0.426 | 0.426 | 0.426 | 0.733 | 0.733 |
| 3EO1 | 0.009 | 0.014 | 0.008 | 0.039 | 0.039 | 0.01 | 0.033 | 0.033 | 0.01 | 0.033 | 0.033 |
| 3EOA | 0.037 | 0.041 | 0.031 | 0.036 | 0.042 | 0.036 | 0.041 | 0.058 | 0.036 | 0.041 | 0.041 |
| 3F1P | 0.012 | 0.047 | 0.147 | 0.466 | 0.466 | 0.012 | 0.479 | 0.579 | 0.012 | 0.06 | 0.479 |
| 3FN1 | 0.011 | 0.011 | 0.034 | 0.49 | 0.49 | 0.013 | 0.442 | 0.442 | 0.011 | 0.442 | 0.442 |
| 3G6D | 0.055 | 0.073 | 0.075 | 0.254 | 0.345 | 0.416 | 0.416 | 0.416 | 0.416 | 0.416 | 0.416 |
| 3H2V | 0.009 | 0.011 | 0.035 | 0.088 | 0.088 | 0.017 | 0.085 | 0.085 | 0.017 | 0.085 | 0.085 |
| 3HI6 | 0.03 | 0.03 | 0.043 | 0.043 | 0.08 | 0.031 | 0.048 | 0.083 | 0.031 | 0.048 | 0.083 |
| 3HMX | 0.004 | 0.005 | 0.026 | 0.038 | 0.256 | 0.038 | 0.202 | 0.202 | 0.038 | 0.202 | 0.202 |
| 3K75 | 0.905 | 0.908 | 0.039 | 0.039 | 0.232 | 0.896 | 0.896 | 0.896 | 0.905 | 0.908 | 0.908 |
| 3L5W | 0.061 | 0.061 | 0.042 | 0.192 | 0.192 | 0.035 | 0.182 | 0.182 | 0.035 | 0.182 | 0.182 |
| 3L89 | 0.008 | 0.011 | 0.008 | 0.019 | 0.019 | 0.015 | 0.015 | 0.019 | 0.015 | 0.015 | 0.019 |
| 3LVK | 0.046 | 0.209 | 0.484 | 0.484 | 0.484 | 0.043 | 0.103 | 0.458 | 0.043 | 0.103 | 0.458 |
| 3MXW | 0.012 | 0.014 | 0.036 | 0.036 | 0.036 | 0.036 | 0.036 | 0.036 | 0.036 | 0.036 | 0.036 |
| 3P57 | 0.02 | 0.024 | 0.087 | 0.087 | 0.087 | 0.02 | 0.07 | 0.07 | 0.02 | 0.07 | 0.07 |
| 3PC8 | 0.786 | 0.812 | 0.824 | 0.824 | 0.824 | 0.869 | 0.869 | 0.869 | 0.786 | 0.869 | 0.869 |
| 3RVW | 0.007 | 0.011 | 0.039 | 0.166 | 0.166 | 0.063 | 0.063 | 0.173 | 0.063 | 0.063 | 0.173 |
| 3S9D | 0.821 | 0.84 | 0.568 | 0.568 | 0.568 | 0.811 | 0.845 | 0.845 | 0.811 | 0.845 | 0.845 |
| 3SGQ | 0.036 | 0.771 | 0.097 | 0.206 | 0.206 | 0.116 | 0.159 | 0.194 | 0.036 | 0.159 | 0.771 |
| 3SZK | 0.01 | 0.011 | 0.063 | 0.063 | 0.063 | 0.009 | 0.063 | 0.063 | 0.009 | 0.063 | 0.063 |
| 3V6Z | 0.006 | 0.008 | 0.031 | 0.076 | 0.076 | 0.014 | 0.052 | 0.074 | 0.014 | 0.052 | 0.074 |
| 3VLB | 0.359 | 0.359 | 0.068 | 0.663 | 0.663 | 0.382 | 0.864 | 0.864 | 0.382 | 0.864 | 0.864 |
| 4CPA | 0.037 | 0.053 | 0.035 | 0.094 | 0.218 | 0.094 | 0.196 | 0.196 | 0.094 | 0.196 | 0.196 |
| 4DN4 | 0.387 | 0.387 | 0.102 | 0.156 | 0.232 | 0.152 | 0.225 | 0.225 | 0.152 | 0.225 | 0.225 |
| 4FZA | 0.62 | 0.678 | 0.01 | 0.048 | 0.048 | 0.01 | 0.015 | 0.042 | 0.62 | 0.678 | 0.678 |
| 4G6J | 0.017 | 0.017 | 0.031 | 0.053 | 0.128 | 0.033 | 0.193 | 0.193 | 0.033 | 0.193 | 0.193 |
| 4G6M | 0.041 | 0.041 | 0.039 | 0.205 | 0.205 | 0.043 | 0.215 | 0.215 | 0.043 | 0.215 | 0.215 |
| 4H03 | 0.004 | 0.072 | 0.016 | 0.022 | 0.03 | 0.021 | 0.03 | 0.03 | 0.021 | 0.03 | 0.03 |
| 4HX3 | 0.023 | 0.036 | 0.014 | 0.065 | 0.065 | 0.014 | 0.058 | 0.063 | 0.014 | 0.058 | 0.063 |
| 4IZ7 | 0.585 | 0.585 | 0.371 | 0.371 | 0.371 | 0.018 | 0.367 | 0.367 | 0.018 | 0.585 | 0.585 |
| 4M76 | 0.008 | 0.008 | 0.009 | 0.052 | 0.052 | 0.03 | 0.03 | 0.042 | 0.03 | 0.03 | 0.042 |
| 7CEI | 0.867 | 0.867 | 0.443 | 0.443 | 0.443 | 0.873 | 0.873 | 0.873 | 0.867 | 0.873 | 0.873 |
| 9QFW | 0.037 | 0.107 | 0.188 | 0.188 | 0.198 | 0.016 | 0.217 | 0.217 | 0.016 | 0.217 | 0.217 |
| BAAD | 0.012 | 0.015 | 0.014 | 0.014 | 0.019 | 0.009 | 0.015 | 0.015 | 0.009 | 0.015 | 0.015 |
| BOYV | 0.014 | 0.014 | 0.035 | 0.037 | 0.037 | 0.016 | 0.038 | 0.038 | 0.016 | 0.038 | 0.038 |
| BP57 | 0.067 | 0.067 | 0.021 | 0.074 | 0.074 | 0.026 | 0.027 | 0.049 | 0.026 | 0.067 | 0.067 |
| CP57 | 0.104 | 0.147 | 0.014 | 0.214 | 0.214 | 0.026 | 0.067 | 0.115 | 0.026 | 0.104 | 0.147 |
| **Average** | 0.269 | 0.316 | 0.127 | 0.219 | 0.263 | 0.240 | 0.326 | 0.356 | 0.266 | 0.372 | 0.416 |

**Table S2:** DockQ score of the best model for the targets in Benchmark 2.

| Target | DockQ score of best model |  |  |  |  |  |  |  |  |  |  |
| --- | --- | --- | --- | --- | --- | --- | --- | --- | --- | --- | --- |
|  | Alphafold |  | Cluspro |  |  | Alphafold refined Cluspro |  |  | Alphafold refined Cluspro plus Alphafold |  |  |
|  | Top 1 | Top 5 | Top 1 | Top 5 | Top 10 | Top 1 | Top 5 | Top 10 | Top 1 | Top 5 | Top 10 |
| 5ZNG | 0.017 | 0.017 | 0.081 | 0.081 | 0.081 | 0.04 | 0.065 | 0.071 | 0.04 | 0.061 | 0.071 |
| 6A6I | 0.151 | 0.151 | 0.103 | 0.514 | 0.514 | 0.106 | 0.191 | 0.191 | 0.106 | 0.179 | 0.191 |
| 6GS2 | 0.486 | 0.52 | 0.004 | 0.076 | 0.438 | 0.487 | 0.487 | 0.487 | 0.486 | 0.51 | 0.52 |
| 6H4B | 0.022 | 0.051 | 0.581 | 0.581 | 0.581 | 0.802 | 0.816 | 0.816 | 0.802 | 0.816 | 0.816 |
| 6IF2 | 0.692 | 0.714 | 0.028 | 0.489 | 0.489 | 0.728 | 0.728 | 0.728 | 0.692 | 0.728 | 0.728 |
| 6II6 | 0.798 | 0.798 | 0.63 | 0.63 | 0.63 | 0.796 | 0.796 | 0.796 | 0.798 | 0.798 | 0.798 |
| 6ONO | 0.619 | 0.619 | 0.02 | 0.089 | 0.089 | 0.639 | 0.649 | 0.649 | 0.639 | 0.649 | 0.649 |
| 6PNQ | 0.17 | 0.17 | 0.044 | 0.053 | 0.212 | 0.041 | 0.22 | 0.22 | 0.041 | 0.22 | 0.22 |
| 6Q76 | 0.925 | 0.925 | 0.019 | 0.598 | 0.598 | 0.944 | 0.944 | 0.944 | 0.925 | 0.944 | 0.944 |
| 6U08 | 0.197 | 0.197 | 0.249 | 0.542 | 0.542 | 0.188 | 0.648 | 0.648 | 0.188 | 0.648 | 0.648 |
| 6ZBK | 0.807 | 0.862 | 0.328 | 0.438 | 0.438 | 0.861 | 0.861 | 0.861 | 0.807 | 0.862 | 0.862 |
| 7AYE | 0.764 | 0.815 | 0.775 | 0.775 | 0.775 | 0.849 | 0.865 | 0.865 | 0.764 | 0.865 | 0.865 |
| 7D2T | 0.006 | 0.01 | 0.46 | 0.46 | 0.46 | 0.013 | 0.489 | 0.489 | 0.013 | 0.489 | 0.489 |
| 7M5F | 0.009 | 0.009 | 0.009 | 0.011 | 0.016 | 0.008 | 0.015 | 0.015 | 0.008 | 0.015 | 0.015 |
| 7N10 | 0.745 | 0.847 | 0.749 | 0.749 | 0.749 | 0.799 | 0.799 | 0.799 | 0.745 | 0.847 | 0.847 |
| 7NLJ | 0.018 | 0.028 | 0.077 | 0.091 | 0.091 | 0.07 | 0.07 | 0.074 | 0.07 | 0.07 | 0.07 |
| 7P8K | 0.032 | 0.061 | 0.022 | 0.478 | 0.478 | 0.886 | 0.886 | 0.886 | 0.886 | 0.886 | 0.886 |
| **Average** | 0.380 | 0.400 | 0.246 | 0.391 | 0.422 | 0.486 | 0.561 | 0.561 | 0.471 | 0.564 | 0.566 |

## Notes

### Competing Interest Statement

The PIPER docking program, used in the ClusPro server, has been licensed by Boston University to Acpharis Inc., and to Schrodinger LLC. The companies sublicense the program for commercial use. However, the ClusPro server is free for academic and governmental research.

### Summary of Updates

Author list updated.

